# Ire1-triggered *hxl1* mRNA splicing coordinates stress tolerance and virulence in the pathogenic fungus *Trichosporon asahii*

**DOI:** 10.64898/2026.06.27.734954

**Authors:** Yuta Shimizu, Yasuhiko Matsumoto, Takashi Sugita

## Abstract

The pathogenic fungus *Trichosporon asahii* causes severe mycoses in immunocompromised hosts, such as neutropenic patients. In *Cryptococcus neoformans*, the unfolded protein response (UPR) sensor Ire1 induces *hxl1* mRNA splicing and contributes to stress responses and virulence. The function of Ire1-triggered *hxl1* mRNA splicing in stress tolerance and virulence of *T. asahii*, however, remains unclear. Here, we demonstrated that *ire1*- and *hxl1* gene-deficient *T. asahii* mutants are sensitive to dithiothreitol (DTT), an inducer of endoplasmic reticulum stress, and exhibit reduced virulence in a silkworm infection model. DTT treatment induced *hxl1* mRNA splicing in the wild-type strain, whereas *ire1* gene-deficient mutants did not undergo *hxl1* mRNA splicing. The *ire1* gene-deficient mutants were more sensitive than the parent strain to DTT, H_2_O_2_, Congo red, and SDS, and showed reduced virulence in silkworms. Similarly, *hxl1* gene-deficient mutants exhibited increased sensitivity to these stressors and reduced virulence. Both the *ire1* gene-deficient and *hxl1* gene-deficient mutants showed decreased expression of reactive oxygen species-detoxifying related genes *CAT2*, *SOD1*, and *SOD2*, compared with the parent strain. Together, these findings suggest that Ire1-triggered *hxl1* mRNA splicing contributes to stress resistance and virulence in *T. asahii*.

## Introduction

*Trichosporon asahii*, a member of the Basidiomycota phylum, is a pathogenic fungus that causes severe invasive infections in immunocompromised hosts, particularly those with neutropenia (1–3). Deep-seated fungal infections caused by *T. asahii* are associated with a high mortality rate (70%–80%) and a poor prognosis (4–9). Echinocandin antifungal agents, commonly used to treat fungal infections, exhibit limited activity against *T. asahii*, and patients receiving micafungin may therefore develop *T. asahii* infections (10, 11). In addition, *T. asahii* strains resistant to the antifungal drugs amphotericin B and azole antifungals have been isolated from infected patients (12, 13). Therefore, *T. asahii* is a clinically significant pathogen, and establishing preventive methods and treatments based on an understanding of its infection mechanisms is critical.

The unfolded protein response (UPR) is involved in adaptation to the host environment and pathogenicity in major fungal pathogens, including *Cryptococcus neoformans*, *Candida albicans*, and *Aspergillus fumigatus* (14–17). Ire1, a key UPR sensor, regulates gene expression by inducing splicing of mRNAs encoding target transcription factors, thereby preventing the accumulation of misfolded proteins (14, 15, 17–19). In these pathogens, Ire1 is involved in endoplasmic reticulum (ER) stress resistance and virulence (14, 15, 20). In *Cryptococcus* species, the stress-responsive transcription factor Hxl1 is activated via Ire1-mediated splicing of *hxl1* mRNA and contributes to ER stress resistance and virulence (15, 16). Although *in silico* analysis has identified a putative *hxl1* gene encoding Hxl1 in *T. asahii* (21), the roles of Ire1 and Hxl1 in stress tolerance and virulence remain unclear. Established gene manipulation methods for *T. asahii,* including the generation of gene-deficient mutants via electroporation and a silkworm infection model for quantitative assessment of virulence have enabled the identification of key virulence factors (22–28).

In this study, we generated *ire1* and *hxl1* gene-deficient *T. asahii* strains and evaluated their responses to various stress conditions and their virulence using a silkworm infection model. We found that dithiothreitol (DTT) induces Ire1-dependent splicing of *hxl1* mRNA in *T. asahii*, and that both Ire1 and Hxl1 play important roles in tolerance to ER and oxidative stress, as well as in virulence in the silkworm infection model.

## Materials & Methods

### Reagents

Nourseothricin was purchased from JENA BIOSCIENCE GMBH (Dortmund, Germany). Hygromycin B and caffeine were purchased from Tokyo Chemical Industry Co., Ltd. (Tokyo, Japan). Cefotaxime sodium, chloroform, 2-propanol, D-glucose, agar, NaCl, sorbitol, H_2_O_2_, and DTT were obtained from FUJIFILM Wako Pure Chemical Corporation (Osaka, Japan). Congo red was purchased from Sigma-Aldrich Co., LLC (St. Louis, MO, USA). G418 was purchased from Enzo Life Science, Inc. (Farmingdale, NY, USA). Hipolypeptone was obtained from NIHON PHARMACEUTICAL Co., LTD (Tokyo, Japan). Tunicamycin (TM) was purchased from Cayman Chemical Company (Ann Arbor, MI, USA). Sodium dodecyl sulfate (SDS) was purchased from NIPPON GENE Co., Ltd. (Tokyo, Japan), and Brefeldin A was obtained from Funakoshi Co., Ltd. (Tokyo, Japan). Nuclease-free water was purchased from Takara Bio Inc. (Shiga, Japan). TRIzol was purchased from Thermo Fisher Scientific Inc. (Waltham, MA, USA).

### T. asahii strains

*T. asahii* MPU129 *ku70*Δwas used as the parent strain (23, 27). Three *ire1* gene-deficient mutants and an *hxl1* gene-deficient mutant were generated from the parent strain. An *hxl1* gene-complemented strain, in which the *hxl1* gene was reintroduced into the *hxl1* gene-deficient mutant, was also constructed. Strains used in this study are listed in Table 1. All experiments involving *T. asahii* were approved by the Biosafety Committee of Meiji Pharmaceutical University. Recombinant DNA experiments were performed in accordance with the approval of the Recombinant DNA Experiment Safety Committee of Meiji Pharmaceutical University (Approval No. 201902).

**Table 1.** Strains used in this study.

| *T. asahii* strains | Relevant genotype | Background | Reference |
| --- | --- | --- | --- |
| MPU129 $\Delta ku70$ (Parent strain) | *ku70::nptII* | MPU129 | Matsumoto Y., *et al.* (23, 27) |
| *ire1*Δ #1-3 | *ku70::nptII, ire1::NAT1* | MPU129 $\Delta ku70$ | This study |
| *hxl1*Δ #1-3 | *ku70::nptII, hxl1::NAT1* | MPU129 $\Delta ku70$ | This study |
| Hxl1 (Complement) | *ku70::nptII, NAT1::hxl1, hph* | MPU129 $\Delta ku70 \Delta hxl1$ #1 | This study |

### *T. asahii* culture conditions

The *T. asahii* used in this study were cultured in SA (Sabouraud agar) medium (1% Hipolypeptone, 4% dextrose, and 1.5% agar) containing G418 (100 µg/mL) and cefotaxime (100 µg/mL) at 27°C for 2 days, as previously described (24, 26).

### Estimation of *T. asahii* Ire1 and Hxl1 by *in silico* analysis

The amino acid sequences of *C. neoformans* Ire1 and Hxl1 were obtained from the National Center for Biotechnology Information (NCBI) database (https://www.ncbi.nlm.nih.gov/). Putative *T. asahii* Ire1 and Hxl1 homologs were identified using BLASTp (https://blast.ncbi.nim.nih.gov/Blast.cgi) based on the corresponding *C. neoformans* sequences. The protein kinase/endoribonuclease with the highest sequence homology was designated Ire1, and the hypothetical protein A1Q2_03745 was designated Hxl1.

### Conservation of Ire1 and Hxl1 amino acid sequences between *T. asahii* and representative fungi

The amino acid sequences of Ire1 proteins from *C. albicans*, *C. neoformans*, and *S. cerevisiae*, as well as the IreA protein from *A. fumigatus*, were obtained from the NCBI database. The *T. asahii* Ire1 sequence was compared with these sequences using BLASTp. Similarly, the sequence of *T. asahii* Hxl1 protein was compared with Hxl1 proteins from *C. gattii*, *C. deuterogattii*, *C. neoformans*H99, and *C. neoformans* JEC21 using BLASTp (https://blast.ncbi.nlm.nih.gov/Blast.cgi). Phylogenetic analysis was performed using the maximum likelihood method implemented in MEGA11 Software (https://www.megasoftware.net/).

### Generation of *T. asahii* gene-deficient mutants and the gene-complemented strain

Gene-deficient and complemented strains were generated as described previously (24, 26, 28). For construction of the *ire1* deletion cassette, approximately 1500–2000 bp of the 5′- and 3’-UTR regions of the *T. asahii ire1* gene were cloned upstream and downstream of the *NAT1* selection marker in the pUC19-NAT1 vector using the In-Fusion HD Cloning Kit (Takara Bio Inc.). The resulting 5′-UTR-NAT1-3′UTR fragment was amplified by polymerase chain reaction (PCR). The *hxl1* deletion cassette was constructed in the same manner using the corresponding 5’- and 3’-UTR regions. Competent cells for electroporation were prepared as described previously (28). Cells of the *T. asahii* parent strain were plated on SA medium and cultured at 27°C for 1 day. Cells were collected from the agar surface, suspended in 2 mL saline, and filtered through a 40-µm pore-size cell strainer (Corning Inc., NY, USA). The filtered suspension was centrifuged at 13,000 rpm for 15 min to obtain a pellet. Cells were resuspended in 10 mL of cold distilled water and transferred to a 1.5-mL tube, followed by centrifugation at 14,000 rpm for 5 min. The pellet was resuspended in 1 mL of cold distilled water containing DTT (final conc. 5 mM) and centrifuged at 14,000 rpm for 5 min. This washing step was repeated three times. The washed cells were resuspended in 1 mL of 1.2 M sorbitol solution, centrifuged at 14,000 rpm for 5 min, and finally resuspended in 100 µL of 1.2 M sorbitol solution to prepare competent cells. For transformation, 2 µL of the PCR-amplified deletion fragment (5’-UTR-NAT1-3’-UTR) was added to 100 µL of competent cells and incubated on ice for 15 min. The mixture was transferred to a 2-mm gap cuvette (Bio-Rad Laboratories, Inc., CA, USA) and electroporated using a Gene Pulser Xcell (Bio-Rad Laboratories, Inc.) under the following conditions: 1800 V, 5 ms (time constant protocol). After electroporation, cells were suspended in 1 mL of YPD (yeast extract peptone dextrose) medium containing 0.6 M sorbitol and cultured at 27°C for 12 h. The cells were then plated onto SA medium containing nourseothricin (300 µg/mL) and incubated at 27°C for 10 days to allow colony formation. Genomic DNA was extracted from the nourseothricin-resistant transformants, and deletion of the *ire1*or *hxl1* gene was confirmed by PCR using the primers listed in Table S3.

Competent cells for electroporation to establish the *hxl1* gene-complemented strain were prepared as described above, with slight modifications. The *hxl1* gene-deficient *T. asahii* mutant was plated on YPD medium and cultured at 27°C for 1 day. Cells were collected from the agar surface, suspended in 3 mL saline, and filtered through a 40-µm cell strainer. The cell suspension was centrifuged at 13,000 rpm for 15 min to obtain a pellet. The pellet was resuspended in 10 mL of chilled distilled water and transferred to a 1.5-mL tube, followed by centrifugation at 14,000 rpm for 5 min. The pellet was then resuspended in 1 mL of cold distilled water and centrifuged at 14,000 rpm for 5 min. The washing step was repeated four times. The washed cells were suspended in 1 mL of 1.2 M sorbitol solution, centrifuged at 14,000 rpm for 5 min, and finally resuspended in 40 µL of 1.2 M sorbitol solution to prepare competent cells.

For transformation, 2 µL of the PCR-amplified DNA fragment [5’-UTR (*hxl1*)-*hxl1*-*hph*-3’-UTR (*hxl1*)] was added to 40 µL of competent cells of the *hxl1* gene-deficient mutant. Electroporation was performed using a time constant protocol (1500 V, 5 ms). After electroporation, cells were suspended in 1 mL of YPD medium containing 0.6 M sorbitol and cultured at 27°C for 12 h. The cells were then plated onto YPD medium containing hygromycin B (200 µg/mL) and cultured at 27°C for 5 days. Genomic DNA was extracted from hygromycin B–resistant transformants, and reintroduction of the *hxl1* and *hph* genes was confirmed by PCR using the primers listed in Table S3.

### Temperature sensitivity test

Temperature sensitivity test was performed as described previously (24, 26, 28). *T. asahii* was cultured on SA medium containing cefotaxime (100 µg/mL) at 27°C for 2 days. Cells were harvested, suspended in saline, and filtered through a 40-µm cell strainer. Cell suspensions were adjusted to an absorbance of 1.0 at 630 nm, and tenfold serial dilutions were prepared in saline. Aliquots (5 µL) of each dilution were spotted onto SA medium. Plates were incubated at 27°C, 37°C, or 40°C, and photographed after 72 h.

For liquid culture assays, Sabouraud liquid medium (1% hipolypeptone, 4% dextrose) was used. Cells were prepared as described above, and the suspension was adjusted to an absorbance 0.005 at 630 nm. Each fungal suspension was incubated at 27°C, 37°C, or 40°C for 96 h. Growth was monitored by measuring absorbance at 630 nm every 24 h using a microplate reader (iMark™ microplate reader; Bio-Rad Laboratories Inc.).

### Chemical stress sensitivity test

Chemical stress sensitivity test was performed as described previously (24, 26, 28). *T. asahii* strains were cultured on SA medium containing cefotaxime (100 µg/mL) at 27°C for 2 days. Cells from each *T. asahii* strain were suspended in saline and adjusted to an absorbance of 1.0 at 630 nm. Tenfold serial dilutions were prepared in saline, and 5 µL of each dilution was spotted onto SA medium containing caffeine, SDS, H_2_O_2_, Congo red, TM, DTT, or Brefeldin A. Plates were incubated at 27°C and photographed after 72 h.

### Silkworm infection experiments

Silkworm infection experiments were performed as described previously (24, 26, 28). Eggs of silkworms (KINSYU × SHOWA) were purchased from Ehime-Sanshu Co, Ltd. (Ehime, Japan). *T. asahii* was cultured in YPD medium containing cefotaxime (100 µg/mL) at 27°C for 2 days. Cells were harvested, suspended in saline, and filtered through a 40-µm cell strainer. The suspension was adjusted to an absorbance of 0.2 at 630 nm, diluted 200-fold with saline, and 50 µL was injected into the dorsal hemolymph of the silkworm using a 1-mL tuberculin syringe (Terumo Medical Corporation, Tokyo, Japan). Silkworms were maintained on an artificial diet (Silkmate 2S), and survival was monitored at 37°C.

The median lethal dose (LD_50_), defined as the number of *T. asahii* cells required to kill 50% of the silkworms, was determined as described previously (24, 26, 28). *T. asahii* (1.1 × 10^1^–9.0 × 10^7^ cells/50 µL) was injected into the hemolymph of silkworms, which were then maintained at 37°C on an artificial diet. The number of surviving silkworms (n = 4/group) was determined after 72 h. The LD_50_ was calculated from dose-response curves obtained by combining the results of three independent experiments using a simple logistic regression model in Prism (GraphPad Software, LLC, San Diego, CA, USA, https://www.graphpad.com/scientific-software/prism/).

### *Hxl1* mRNA splicing assay

*T. asahii* was cultured on SDA containing cefotaxime (100 µg/mL) for 2 days. Cells from each strain were suspended in saline and filtered through a 40-µm cell strainer. The suspension was adjusted to an absorbance of 2.0 at 600 nm, centrifuged at 14,000 rpm for 5 min, and the supernatant was removed. Sabouraud dextrose liquid medium (500 µL) with or without DTT (final conc. 48 mM) was added to each cell pellet. After incubation at 27°C for 0, 10, 20, or 30 min, cells were centrifuged at 14,000 rpm for 5 min, and the supernatant was discarded. TRIzol (1 mL) was added to each pellet, and the cells were resuspended. The suspension was transferred to a ZR Bashing Bead Lysis tube and vortexed at 1000 rpm for 5 min. Chloroform (200 µL) was added, and the mixture was incubated at room temperature for 5 min, followed by centrifugation at 12,000 x g for 15 min. The aqueous phase (500 µL) was transferred to an EP tube, and 2-propanol (500 µL) was added. After incubation at room temperature for 10 min, the mixture was centrifuged at 12,000 x g for 10 min, and the supernatant was discarded. The RNA pellet was resuspended in 60 µL of nuclease-free water. Reverse transcription (RT) was performed using ReverTra Ace qPCR RT Master Mix (TOYOBO Co., Ltd., Osaka, Japan). Band intensities of unspliced and spliced RT-PCR products were quantified using Image J (https://imagej.net/ij/download.html) and analyzed based on the combined results of three independent experiments. The primers used for RT-PCR are listed in Table S3.

### Gene expression analysis using quantitative RT-PCR

*T. asahii* was cultured on SDA containing cefotaxime (100 µg/mL) for 2 days. Cells from each strain were suspended in saline and filtered through a 40-µm cell strainer. The suspension was adjusted to an absorbance of 2.0 at 600 nm. The suspension was centrifuged at 14,000 rpm for 5 min, and the supernatant was removed. Sabouraud dextrose liquid medium (500 µL) with or without DTT (final conc. 3 mM) was added to each cell pellet. After incubation at 27°C for 30 min, cells were centrifuged at 14,000 rpm for 5 min, and the supernatant was discarded. TRIzol solution (1 mL) was added to each cell pellet, and the cells were resuspended. The suspension was transferred to a ZR Bashing Bead Lysis tube and vortexed at 1000 rpm for 5 min. Chloroform (200 µL) was added, and the mixture was incubated at room temperature for 5 min, followed by centrifugation at 12,000 x g for 15 min. The aqueous phase (500 µL) was transferred to an EP tube and 2-propanol (500 µL) was added. After incubation at room temperature for 10 min, the mixture was centrifuged at 12,000 x g for 10 min, and the supernatant was discarded. The RNA pellet was resuspended in 60µL of nuclease-free water. Reverse transcription was performed using ReverTra Ace qPCR RT Master Mix. Quantitative RT-PCR was performed using KOD SYBR qPCR mix (TOYOBO Co., Ltd., Osaka, Japan) on a Mygo Mini system (Funakoshi Co., Ltd., Tokyo, Japan). *ACT1* and *GAPDH* were used as endogenous controls. The primers used in this study are listed in Table S3.

## Statistical analysis

All experiments were performed at least three times, and representative results are shown. For *T. asahii* growth assays, statistical significance was calculated using Student’s *t*-test. For silkworm infection experiments, survival curves were generated using the Kaplan-Meier method, and differences between groups were assessed using the log-rank test in JMP Pro 17 (https://www.jmp.com/ja_jp/support/jmp-software-updates.html). A P value < 0.05 was considered statistically significant. For quantitative RT-PCR analyses, data are presented as mean ± standard error, and statistical significance was calculated using Student’s *t*-test.

## Data availability

Data will be made available upon reasonable request.

## Results

### Ire1-triggered *hxl1* mRNA splicing in *T. asahii* under UPR stress

A putative Ire1-triggered splicing site within the *hxl1* mRNA sequence of *T. asahii* was identified by *in silico* analysis (21) (Fig. 1A). In *C. neoformans*, DTT, an inducer of UPR stress, promotes *hxl1* mRNA splicing (15). Therefore, to determine whether DTT treatment promotes *hxl1* mRNA splicing in *T. asahii*, RT-PCR was performed using the primers shown in Fig. 1B. DTT treatment increased the level of spliced *hxl1* mRNA in a time-dependent manner and decreased the amount of unspliced *hxl1* mRNA (Fig. 1C, D). Similarly, spliced *hxl1* mRNA levels in *T. asahii* increased in a DTT concentration-dependent manner, while unspliced *hxl1* mRNA levels decreased (Fig. 1E, F). These findings indicate that DTT treatment promotes *hxl1* mRNA splicing in *T. asahii*.

**Fig. 1.**
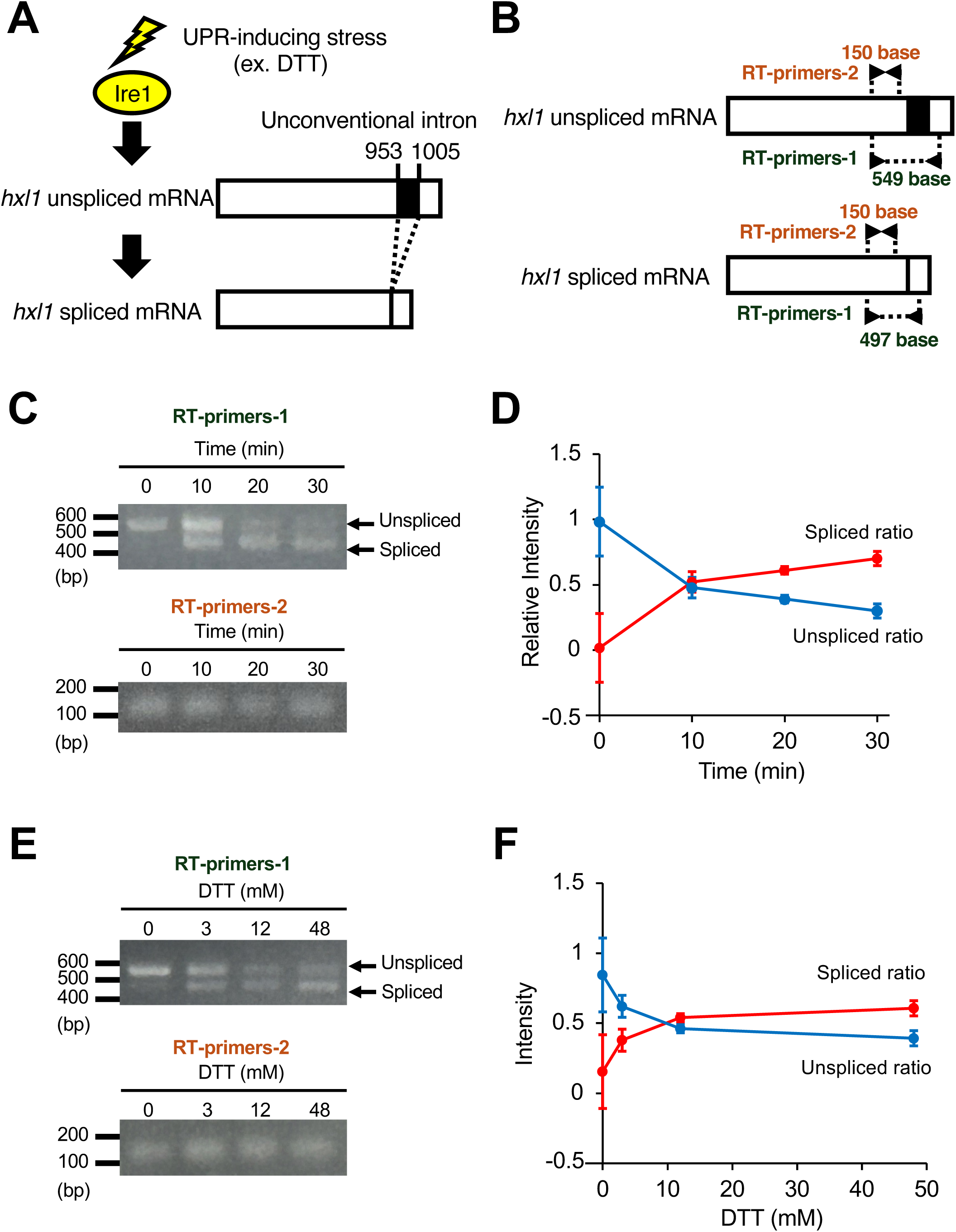
Splicing of *hxl1* mRNA by Ire1 in *T. asahii* by DTT treatment. (**A**) Schematic illustration of *hxl1* mRNA splicing by Ire1 in *T. asahii*. (**B**) Splicing site and primers used in RT-PCR. (**C**) Time-course analysis of *hxl1* mRNA splicing following DTT treatment. The *T. asahii* parent strain was treated with dithiothreitol (DTT; 48 mM) for 0, 10, 20, or 30 min. RT-PCR was performed to detect *hxl1* mRNA splicing, and PCR products were analyzed by DNA gel electrophoresis. (**D**) Quantification of spliced *hxl1* mRNA (red) and unspliced *hxl1* mRNA (blue) in the time-course experiment. Relative ratios were calculated. n = 3/group. (**E**) Dose-response analysis of *hxl1* mRNA splicing following DTT treatment. The parent strain was treated with DTT (0, 3, 12, 48 mM) for 30 min. RT-PCR was performed as described above, and PCR products were analyzed by DNA gel electrophoresis. (**F**) Quantification of spliced *hxl1* mRNA (red) and unspliced *hxl1* mRNA (blue) in the dose-response experiment. Relative ratios were calculated. N = 3/group.

### Conservation of the *T. asahii* Ire1 protein in representative fungi

We performed amino acid sequence homology analysis of the *T. asahii* Ire1 protein using sequences from *S. cerevisiae*, a representative fungus, and the pathogenic fungi *C. neoformans*, *C. albicans*, and *A. fumigatus*. The *T. asahii* Ire1 protein shared 38%, 50%, 41%, and 32% amino acid identity with the Ire1 proteins of *S. cerevisiae*, *C. neoformans*, *C. albicans*, and *A. fumigatus*, respectively (Table S1). Phylogenetic analysis indicated that *T. asahii* Ire1 protein is most closely related to the *C. neoformans* Ire1 protein among the species examined (Fig. 2A). The *T. asahii* Ire1 protein contains functional domains Luminal_IRE1, STKc_IRE1, and RNase_Ire1, similar to the Ire1 proteins in other fungi (Fig. 2B). In addition, key functional regions, including the homodimer interface within the Luminal_IRE1 domain, the ATP binding site within the STKc_IRE1 domain, and the kinase and homodimer interfaces within the RNase_Ire1 domain, were conserved across all fungi analyzed (Fig. 2C).

**Fig. 2.**
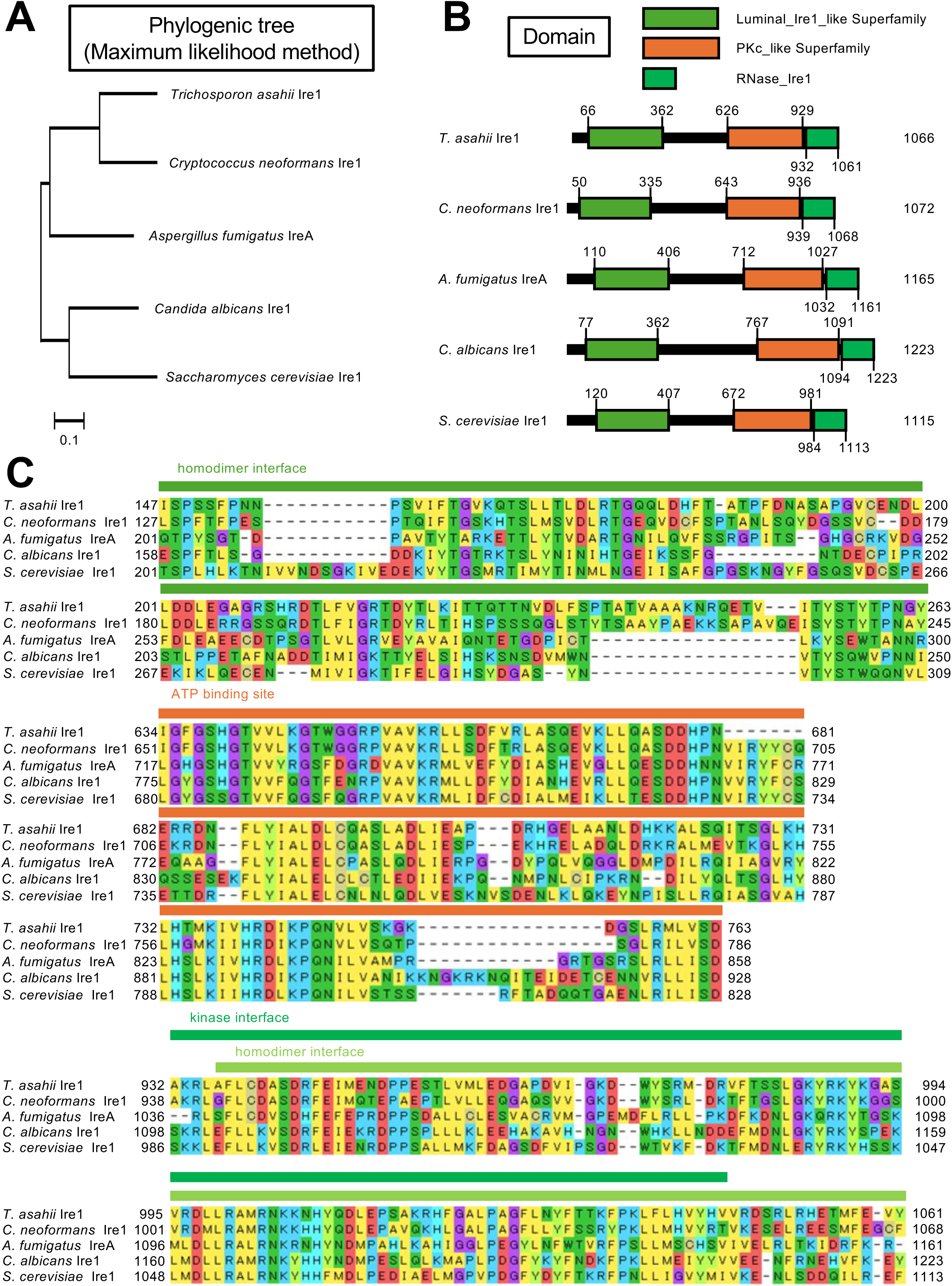
Conservation of the *T. asahii* Ire1 protein among representative fungi. (**A**) Phylogenetic tree of Ire1 proteins from *T. asahii*, *C. neoformans*, *C. albicans*, and *S. cerevisiae*, and the IreA protein of *A. fumigatus*, constructed using the maximum likelihood method. (**B**) Domain architecture of the Ire1 and IreA proteins. (**C**) Amino acid sequences of the Luminal_IRE1, STKc_IRE1, and RNase_Ire1 domains of the Ire1 and IreA proteins. MEGA11 Software was used for sequence alignment analysis and domain prediction.

### Generation of the *ire1* gene-deficient *T. asahii* mutants

Using an established method for generating gene-deficient *T. asahii* strains using homologous recombination (23, 25, 27), the *ire1* gene was replaced with the *NAT1* gene, which confers nourseothricin resistance, thereby generating *ire1* gene-deficient mutants (Fig. 3A). Transformants resistant to nourseothricin were obtained by introducing a DNA fragment containing the *NAT1* gene flanked by the 5’- and 3’-UTRs of *ire1* into *T. asahii* cells via electroporation (Fig. 3B). Genomic DNA was extracted from three independent transformant strains, and deletion of the *ire1* gene was confirmed by PCR (Fig. 3C, D). These results suggest that three independent *ire1* gene-deficient mutants (*ire1*Δ#1–3) were successfully generated (*ire1*Δ#1–3).

**Fig. 3.**
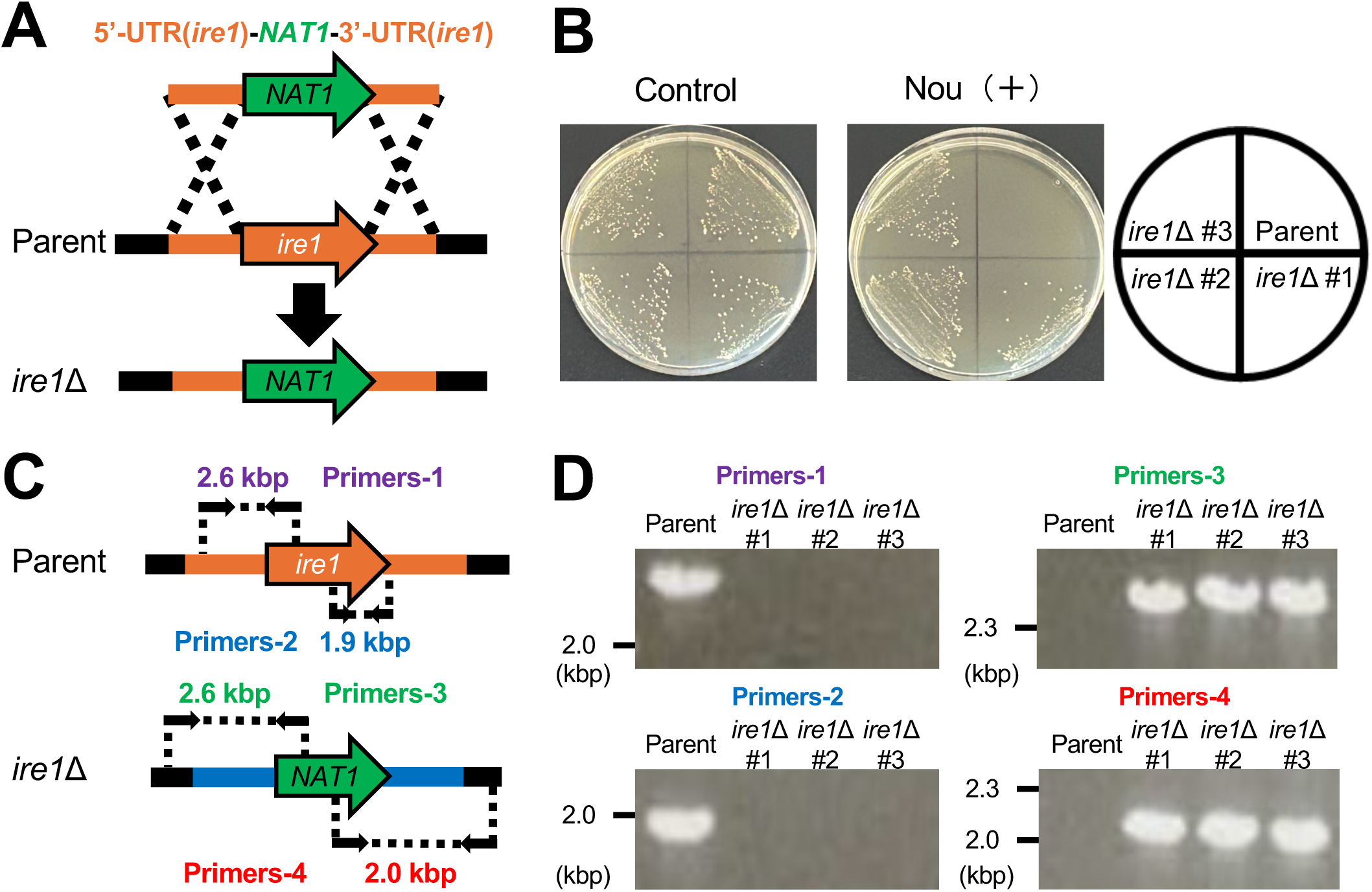
Generation of *ire1* gene-deficient *T. asahii* mutants. (**A**) Schematic illustration of the strategy used to generate the *ire1* gene-deficient mutants. The genome structure of the deletion mutant is shown. (**B**) The parent strain (Parent) and *ire1* gene-deficient mutants (*ire1*Δ #1-3) were plated on SA medium containing nourseothricin (Nou; 100 µg/mL) and cultured at 27°C for 2 days. (**C**) Positions of primers and expected PCR product sizes for the parent and *ire1* gene-deficient mutants. (**D**) DNA gel electrophoresis of PCR products used to confirm deletion of the *ire1* gene in the mutants (*ire1*Δ #1-3).

### Role of the *ire1* gene in *hxl1* mRNA splicing in *T. asahii*

In *C. neoformans*, Ire1 is involved in the splicing of its target *hxl1* mRNA (15). We therefore investigated whether *ire1* is required for DTT-induced *hxl1* mRNA splicing in *T. asahii*. In the parent strain, DTT treatment increased the levels of spliced *hxl1* mRNA, whereas no such increase was observed in *ire1* gene-deficient mutants (Fig. 4). These results suggest that *ire1* is required for DTT-induced *hxl1* mRNA splicing in *T. asahii*.

**Fig. 4.**
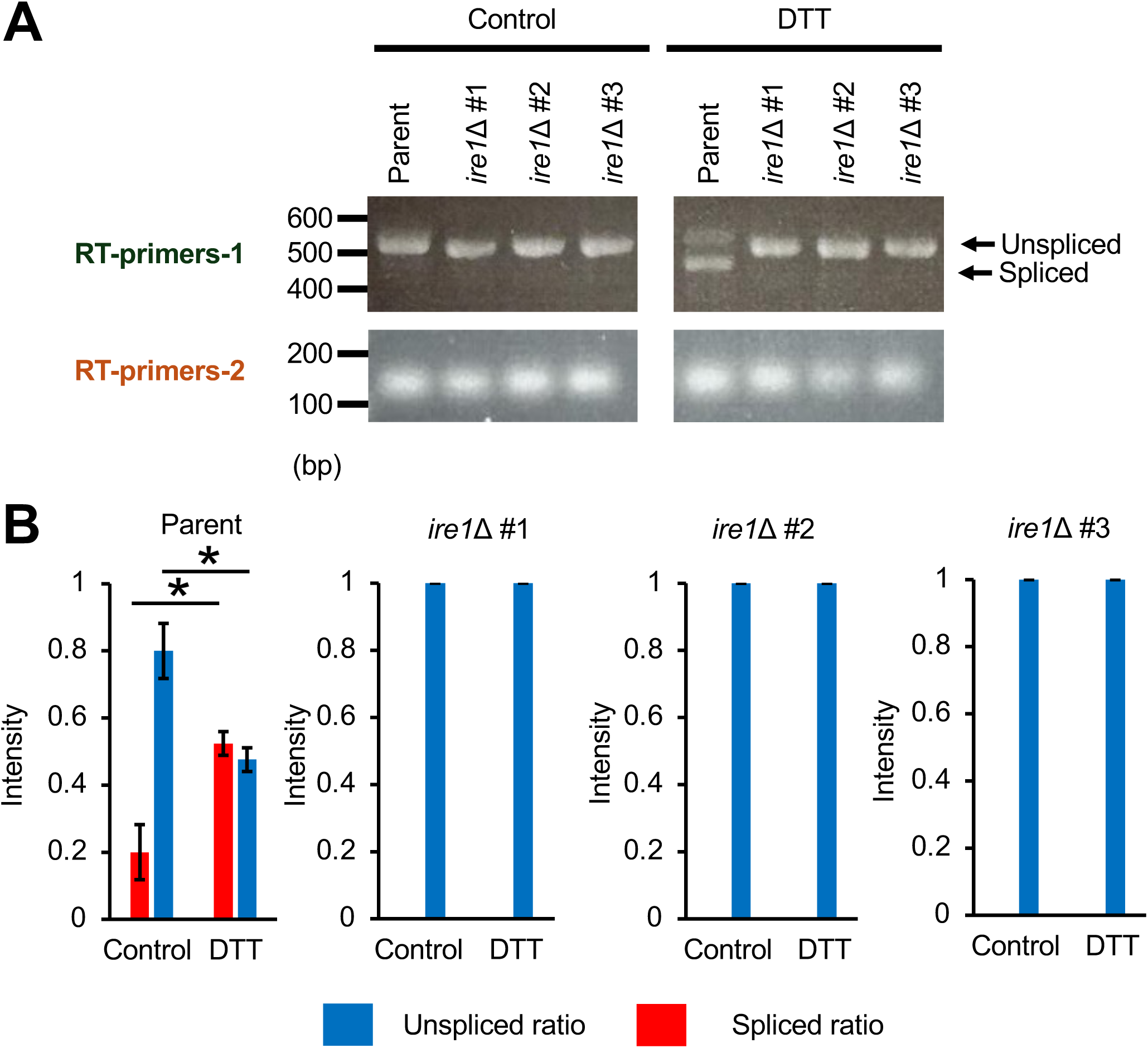
**Requirement of *ire1* for *hxl1* mRNA splicing in *T. asahii*** (**A**) The *T. asahii* parent strain (Parent) and the *ire1* gene-deficient mutants (*ire1*Δ #1-3) were treated with DTT (48 mM) for 30 min. RT-PCR was performed to detect *hxl1* mRNA splicing, and PCR products were analyzed by DNA gel electrophoresis. (**B**) Quantification of spliced *hxl1* mRNA (red) and unspliced *hxl1* mRNA (blue) in the time course experiment. Relative ratios were calculated. n = 3/group. *: P < 0.05.

### Role of *ire1* in *T. asahii* stress tolerance

In *C. neoformans*, *ire1* gene-deficient mutants exhibit sensitivity to 37°C, ER stress, oxidative stress, cell wall stress, and cell membrane stress (15, 16, 29, 30). Similarly, *C. albicans ire1* gene-deficient mutants are sensitive to ER stress, cell wall stress, and cell membrane stress (17, 20, 31), and *S. cerevisiae ire1* gene-deficient mutants exhibit sensitivity to ER stress (18, 32). Therefore, we investigated whether *ire1* contributes to stress tolerance in *T. asahii*. The *T. asahii ire1* gene-deficient mutants exhibited delayed growth at 27°C and 37°C relative to the parent strain, with significant growth delay at 40°C (Fig. 5A, B). Moreover, the mutants exhibited delayed growth on SDA containing DTT, H₂O₂, Congo red, or SDS (Fig. 5C). On the other hand, no marked sensitivity was observed under treatment with TM, caffeine, or Brefeldin A (Fig. 5C). These findings indicate that *ire1 T. asahii* is involved in growth and tolerance to high-temperature stress, ER stress, oxidative stress, and cell wall and membrane stress.

**Fig. 5.**
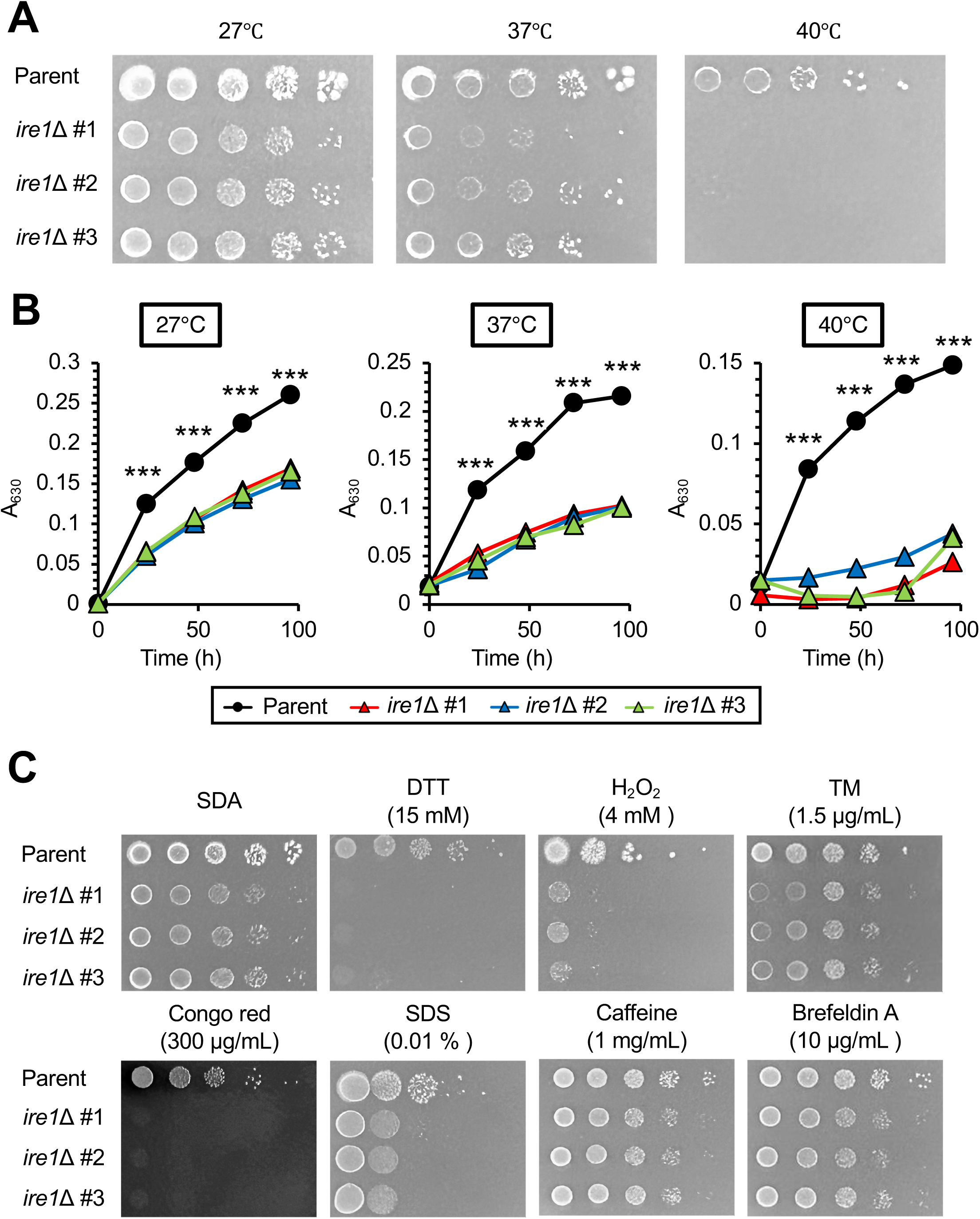
Stress sensitivity of *ire1* gene-deficient *T. asahii* mutants. (**A**) The parent strain (Parent) and *ire1* gene-deficient mutants (*ire1*Δ #1-3) were cultured on SA medium at 27°C for 2 days. *T. asahii* cells were suspended in saline and serially diluted 10 times. Aliquots (5 μL) were spotted onto SA medium and incubated at 27°C, 37°C, or 40°C for 72 h. (**B**) The parent strain (Parent) and *ire1* gene-deficient mutants (*ire1*Δ #1-3) were cultured in Sabouraud dextrose liquid medium at 27°C, 37°C, or 40°C, and growth was monitored by measuring absorbance at 630 nm (**C**) Cell suspensions were spotted onto SA medium containing dithiothreitol (DTT; 15 mM), H_2_O_2_ (4 mM), tunicamycin (TM; 1.5 µg/mL), Congo red (300 µg/mL), sodium dodecyl sulfate (SDS; 0.01%), caffeine (1 mg/mL), or Brefeldin A (10 µg/mL), and incubated at 27°C for 72 h. *: P < 0.05. ***: P < 0.001.

### Decreased virulence of *T. asahii ire1* gene-deficient mutants in silkworms

In *C. neoformans*, *C. albicans*, and *A. fumigatus*, Ire1 contributes to pathogenicity in mice (14–17). An established silkworm infection model was used to evaluate the virulence of *T. asahii* (22–24, 26, 28). We next investigated whether *ire1* is involved in the virulence of *T. asahii* in the silkworm model. Silkworms infected with *ire1* gene-deficient mutants exhibited prolonged survival compared to those infected with the parent strain (Fig. 6A). To quantitatively assess virulence, the LD_50_ value was determined (23, 27). The LD_50_ of the parent strain was 5.0 × 10² cells, whereas the LD_50_ values of the *ire1* gene-deficient mutants exceeded 4.5 × 10⁷ cells (Fig. 6B). These results suggest that *ire1* is required for the virulence of *T. asahii* in the silkworm model.

**Fig. 6.**
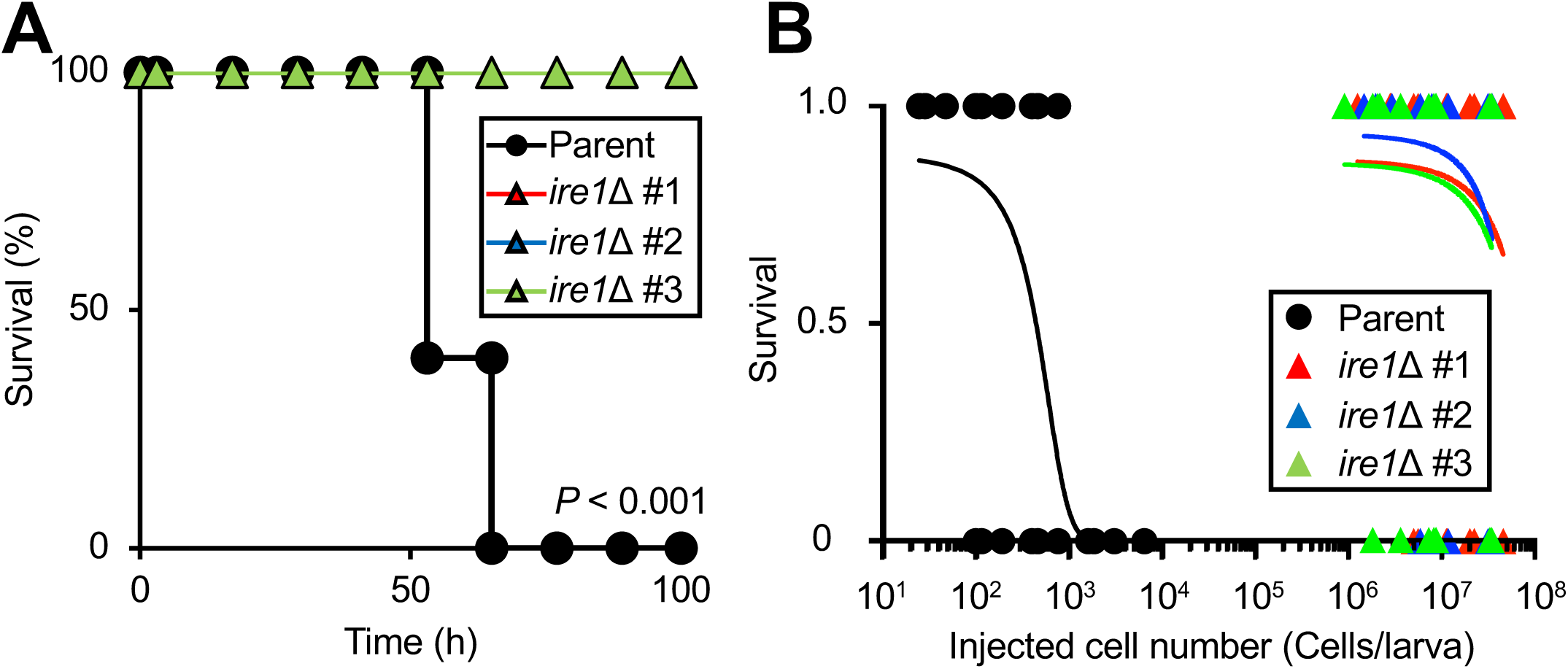
Reduced virulence of *ire1* gene-deficient *T. asahii* mutants in the silkworm infection model. (**A**) The *T. asahii* parent strain (1.6 x 10^3^ cells/larva) and *ire1* gene-deficient mutants (*ire1*Δ #1; 3.5 x 10^3^ cells/larva, *ire1*Δ #2; 2.4 x 10^3^ cells/larva, *ire1*Δ #3; 1.6 x 10^3^ cells/larva) were injected into the silkworm hemolymph, and silkworm survival was monitored for 96 h at 37°C. Differences between the parent strain and mutant groups were assessed using the log-rank test based on Kaplan-Meier survival curves. *: P < 0.05. n = 10/group. (**B**) Dose-response analysis of *T. asahii* infection. The number of surviving silkworms at 37°C was determined 72 h after injection. Survival status was recorded as 1 (alive) or 0 (dead). n = 40–48/group. *: P < 0.05.

### Conservation of the *T. asahii* Hxl1 protein in representative fungi

In *C. neoformans*, Hxl1, a target of Ire1, is involved in stress resistance and virulence (15, 16, 29, 30). Because *T. asahii* belongs to the Basidiomycota phylum, which includes *C. neoformans*, it is possible that the *hxl1* similarly contributes to stress tolerance and virulence in *T. asahii*. We performed amino acid sequence homology analysis of the *T. asahii* Hxl1 protein using sequences from *C. neoformans* H99, *C. neoformans* JEC21, *C. gattii,* and *C. deuterogattii*. The *T. asahii* Hxl1 protein shared 80%, 73%, 31%, and 32% amino acid identity with the Hxl1 proteins of *C. neoformans* H99, *C. neoformans* JEC21, *C. gattii*, and *C. deuterogattii*, respectively (Table S2). Phylogenetic analysis based on amino acid sequences indicated that the *T. asahii* Hxl1 protein is most closely related to the Hxl1 protein of *C. gattii* among the species examined (Fig. 7A). The *T. asahii* Hxl1 protein contains a basic leucine zipper (bZIP) DNA-binding domain, similar to Hxl1 proteins in other fungi (Fig. 7B). Furthermore, the DNA-binding site within the bZIP domain was conserved in all fungi analyzed (Fig. 7C).

**Fig. 7.**
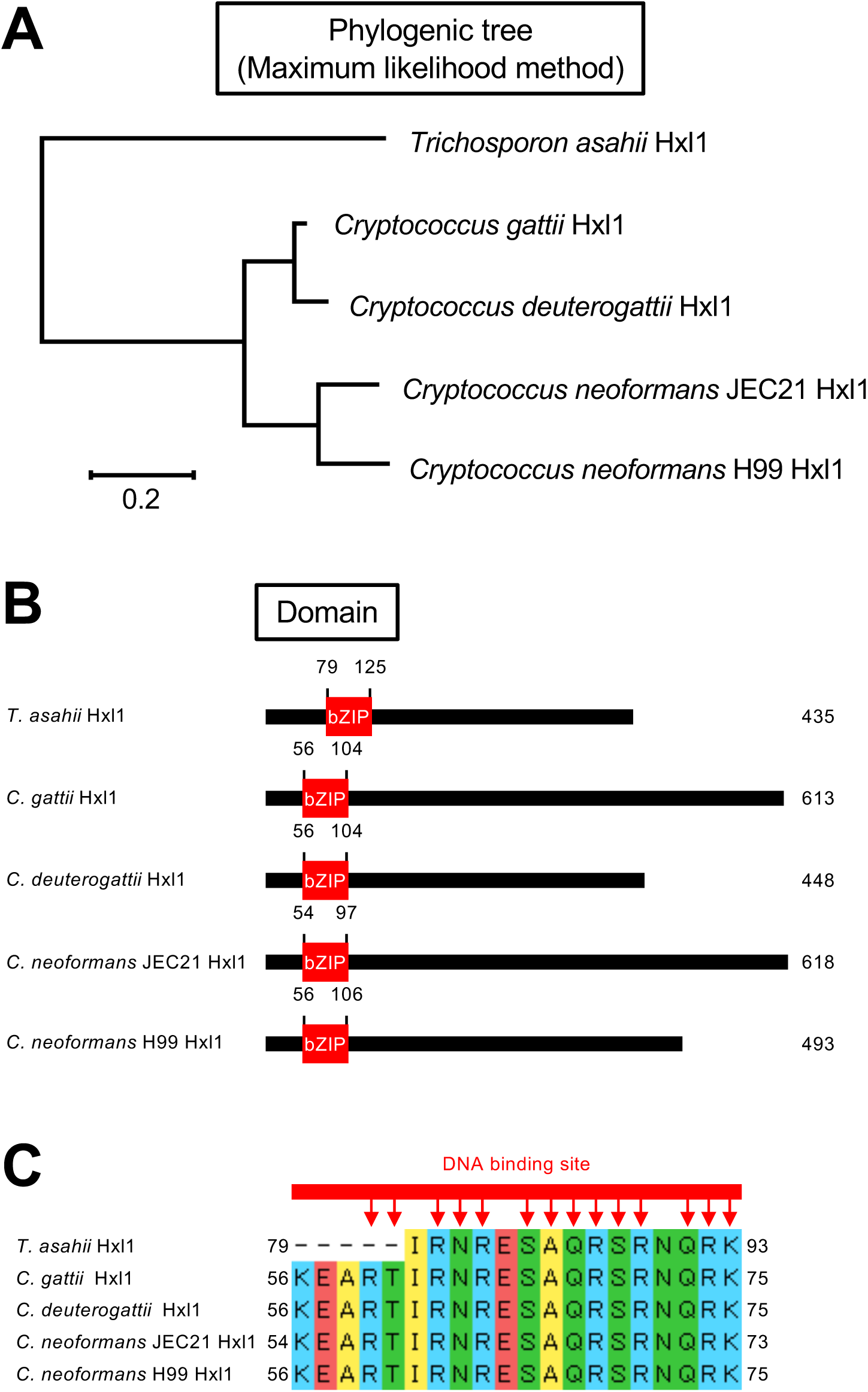
Conservation of the *T. asahii* Hxl1 protein among representative fungi. (**A**) Phylogenetic tree of Hxl1 proteins from *T. asahii, C. gattii*, *C. deuterogattii*, and *C. neoformans* constructed using the maximum likelihood method. (**B**) Domain architecture of Hxl1 proteins. Fungal Hxl1 proteins contain a bZIP_u3 domain. (**C**) Amino acid sequences within the bZIP_u3 domain of Hxl1 proteins. The bZIP_u3 domain includes a region corresponding to the DNA-binding domain. MEGA11 Software was used for sequence alignment and domain prediction.

### Generation of *hxl1* gene-deficient and complemented *T. asahii* strains

An *hxl1* gene-deficient mutant was generated using the same method as that used for *ire1* gene-deficient mutants. The *hxl1* gene was deleted by homologous recombination using targeting DNA fragments (Fig. 8A). Nourseothricin-resistant transformants were obtained by introducing a DNA fragment containing the *NAT1* gene, flanked by the 5’UTR and 3’UTR of *hxl1,* into *T. asahii* cells via electroporation (Fig. 8B). Deletion of the *hxl1* gene in the transformants was confirmed by PCR (Fig. 8C, D). A complemented strain was then generated by reintroducing the *hxl1* gene together with the *hph* gene, which confers hygromycin B resistance, into the *hxl1* gene-deficient mutant via homologous recombination (Fig. 8). Furthermore, RT-PCR analysis using primers designed to detect DTT-induced splicing of *hxl1* mRNA (Fig. 1) showed that the corresponding product detected in the parent strain was absent in the *hxl1* gene-deficient mutant but restored in the complemented strain (Fig. 8E). These results indicate that the *hxl1* gene-deficient and complemented strains were successfully generated.

**Fig. 8.**
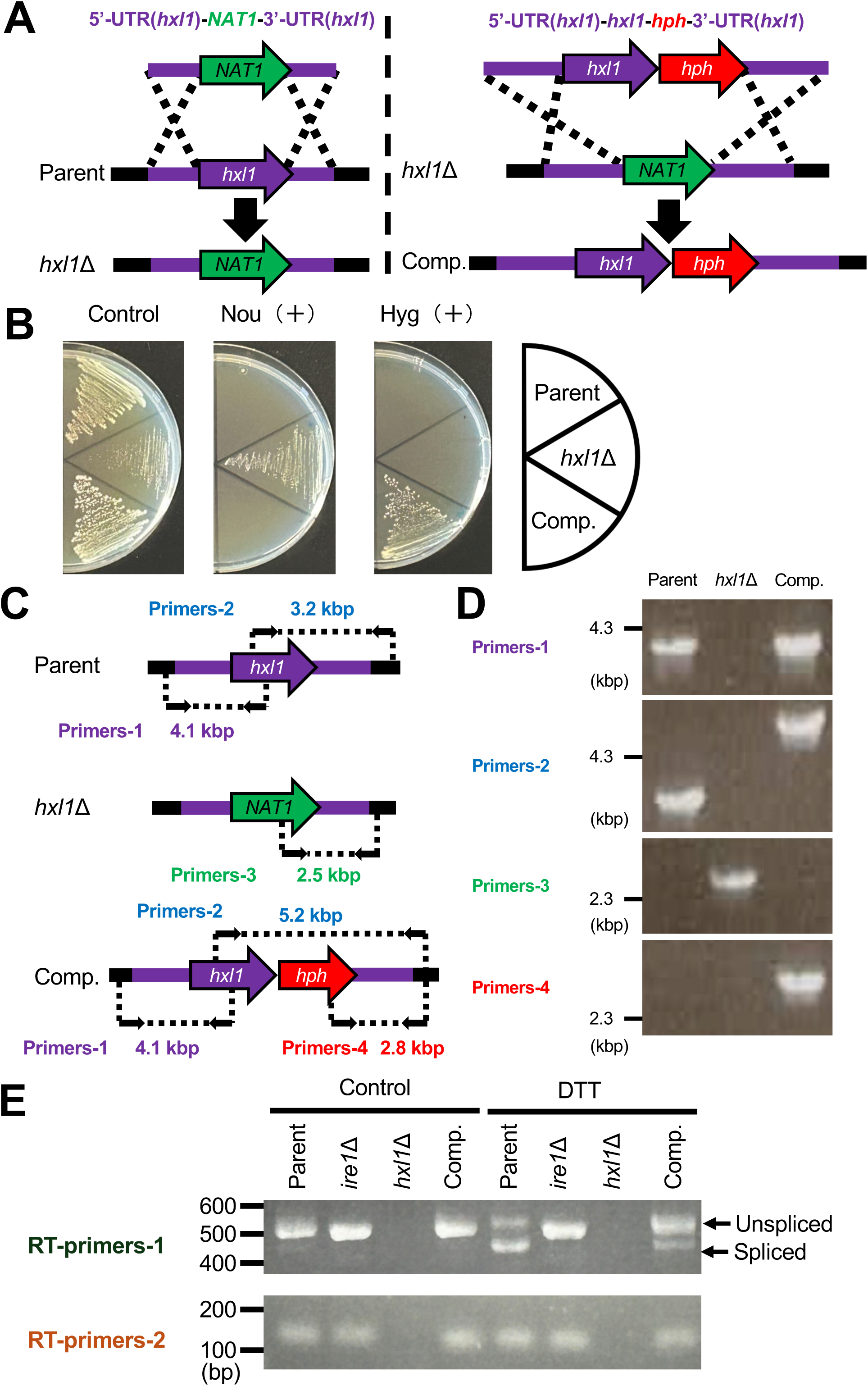
Generation of *hxl1* gene-deficient mutant and-complemented *T. asahii* strains. (**A**) Schematic illustration of the strategy used to generate the *hxl1* gene-deficient mutant and *hxl1* gene-complemented strain. (**B**) The parent strain (Parent), *hxl1* gene-deficient mutant (*hxl1*Δ), and *hxl1* gene-complemented strain (Comp.) were plated on SA medium containing nourseothricin (Nou; 100 µg/mL) or hygromycin B (Hyg; 75 µg/mL) and cultured at 27°C for 2 days. (**C**) Positions of primers and expected PCR product sizes of the parent, *hxl1* gene-deficient mutant, and *hxl1* gene-complemented strain. (**D**) DNA gel electrophoresis of PCR products used to confirm deletion of *hxl1* (*hxl1*Δ) and its reintroduction in the complemented strain (Comp.). (**E**) The parent strain, *ire1* gene-deficient mutant, *hxl1* gene-deficient mutant, and the *hxl1* gene-complemented strain were treated with or without DTT (48 mM) for 30 min. *hxl1* and *act1* mRNA levels were analyzed by RT-PCR, and PCR products were examined by DNA gel electrophoresis.

### Role of *hxl1* in stress tolerance in *T. asahii*

The *T. asahii ire1* gene-deficient mutants exhibited delayed growth at 40°C and sensitivity to DTT, H₂O₂, Congo red, and SDS. Therefore, we investigated whether *hxl1* is involved in stress tolerance in *T. asahii*. The *hxl1* gene-deficient mutant exhibited delayed growth at 27°C and 37°C compared with the parent strain, with more pronounced at 40°C (Fig. 9A, B). In addition, the mutant exhibited delayed growth on SDA containing DTT, H₂O₂, Congo red, and SDS (Fig. 9C). These phenotypes were restored in the *hxl1* gene-complemented strain (Fig. 9). These results suggest that *hxl1* contributes to tolerance to heat stress, ER stress, oxidative stress, and cell wall and membrane stress in *T. asahii*.

**Fig. 9.**
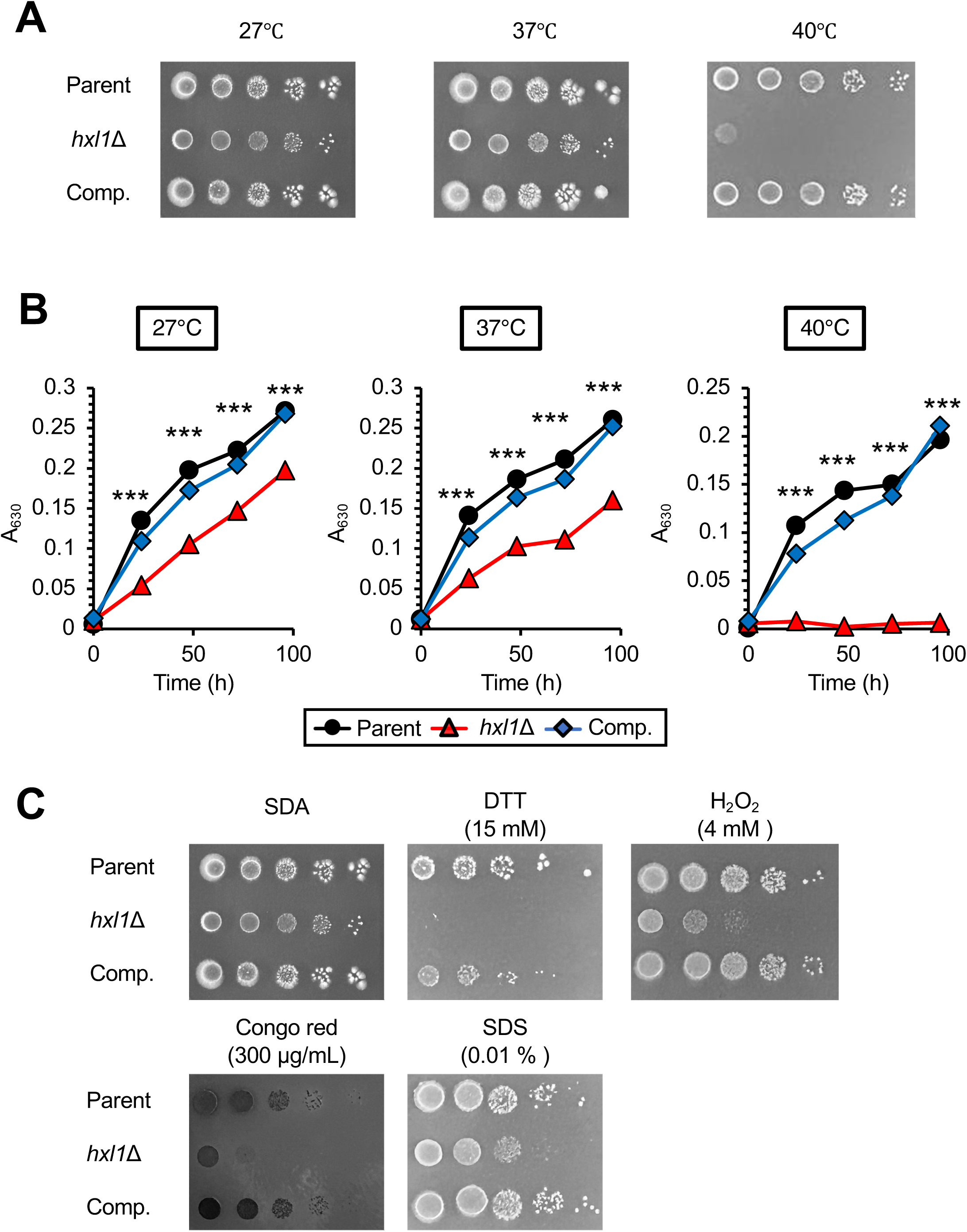
**Stress sensitivity of *hxl1* gene-deficient *T. asahii mutants*** (**A**) The parent strain (Parent), *hxl1* gene-deficient mutant (*hxl1*Δ), and *hxl1* gene-complemented strain (Comp.) were cultured on SA medium at 27°C for 2 days. *T. asahii* cells were suspended in saline and serially diluted 10 times. Aliquots (5 μL) were spotted onto SA medium and incubated at 27°C, 37°C, or 40°C for 72 h. (**B**) The parent strain (Parent), *hxl1* gene-deficient mutant (*hxl1*Δ), and *hxl1* gene-complemented strain (Comp.) were cultured in Sabouraud dextrose liquid medium at 27°C, 37°C, or 40°C, and growth was monitored by measuring absorbance at 630 nm. (**C**) Cell suspensions were spotted onto SA medium containing dithiothreitol (DTT; 15 mM), H_2_O_2_ (4 mM), Congo red (300 µg/mL), and sodium dodecyl sulfate (SDS; 0.01%) and incubated at 27°C for 72 h. *: P<0.05. ***: P<0.001.

### Reduced virulence of *hxl1* gene-deficient mutants in silkworms

The *T. asahii ire1* gene-deficient mutants exhibited reduced virulence in the silkworm model. We therefore investigated whether *hxl1* is also involved in virulence. Silkworms infected with the *hxl1* gene-deficient mutant exhibited prolonged survival compared with those infected with the parent strain (Fig. 10A). The LD_50_ of the *hxl1* gene-deficient mutant was higher than that of the parent strain (Fig. 10B). This phenotype was restored in the *hxl1*-complemented strain (Fig. 10). These results indicate that *hxl1* is required for *T. asahii* virulence in the silkworm infection model.

**Fig. 10.**
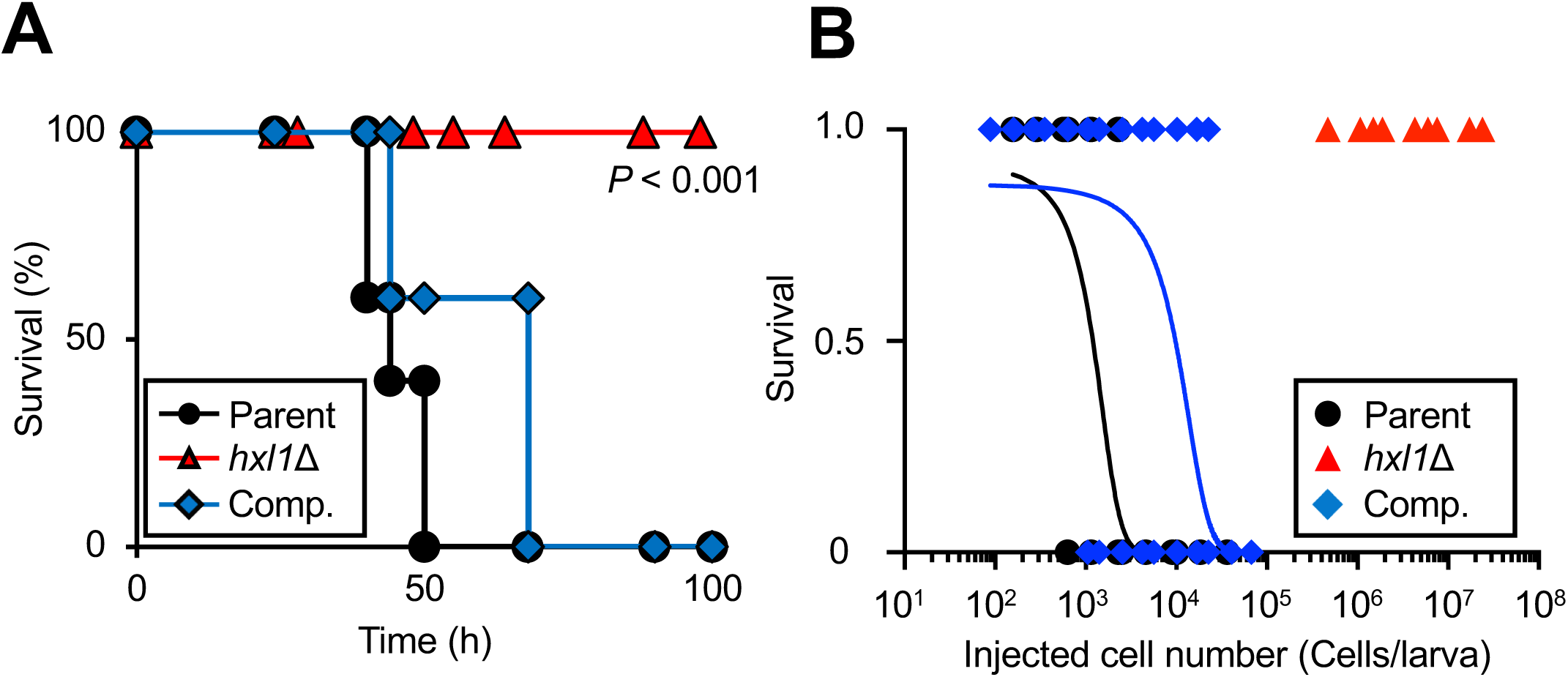
Reduced virulence of *hxl1* gene-deficient *T. asahii* mutants in the silkworm infection model. (**A**) The *T. asahii* parent strain (Parent: 3.1 x 10^3^ cells/larva), *hxl1* gene-deficient mutant (*hxl1*Δ: 4.5 x 10^3^ cells/larva), and *hxl1*-complemented strain (Comp.: 3.9 x 10^3^ cells/larva) were injected into the silkworm hemolymph, and silkworm survival was monitored for 96 h at 37°C. Differences between the parent strain and mutant groups were assessed using the log-rank test based on Kaplan-Meier survival curves. *: P < 0.05. n = 10/group. (**B**) Dose-response analysis of *T. asahii* infection. The number of surviving silkworms at 37°C was determined 72 h after injection. Survival status was recorded as 1 (alive) or 0 (dead). n = 36-84/group. *: P < 0.05.

### Comparison of phenotypes of *T. asahii ire1-* and *hxl1* gene-deficient mutants

Next, we compared the phenotypes of *T. asahii ire1-* and *hxl1* gene-deficient mutants. The delayed growth in the *ire1* gene-deficient mutant at 27°C, 37°C, and 40°C was comparable to that of the *hxl1* gene-deficient mutant (Fig. 11A, B). The *hxl1* gene-deficient mutant exhibited similar sensitivity to DTT and Congo red but was less sensitive to H₂O₂ and SDS than the *ire1* gene-deficient mutant (Fig. 11C). In addition, the *hxl1* gene-deficient mutant exhibited reduced virulence in silkworms, similar to the *ire1* gene-deficient mutant (Fig. 11D, E). These results suggest that although the overall phenotypes are similar, the *hxl1* gene-deficient mutant exhibits slightly weaker sensitivity to oxidative and membrane stress than the *ire1* gene-deficient mutant.

**Fig. 11.**
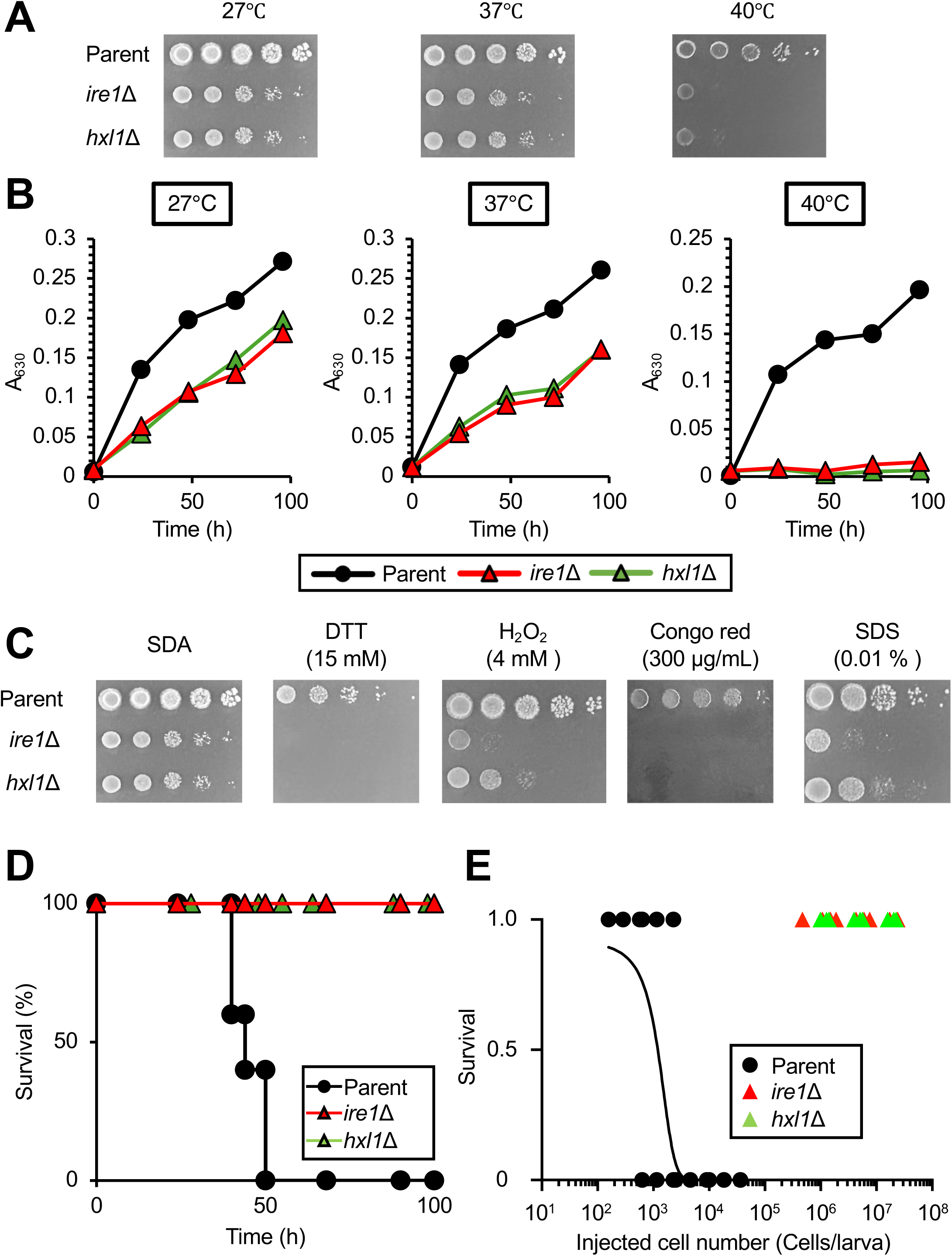
**Phenotypic comparison of *ire1* gene-deficient and *hxl1* gene-deficient *T. asahii***mutants. (**A**) The parent strain (Parent), *ire1* gene-deficient mutant (*ire1*Δ), and *hxl1* gene-deficient mutant (*hxl1*Δ) were cultured on SA medium at 27°C for 2 days. *T. asahii* cells were suspended in saline and serially diluted 10 times. Aliquots (5 μL) were spotted onto SA medium and incubated at 27°C, 37°C, or 40°C for 72 h. (**B**) The parent strain (Parent), *ire1* gene-deficient mutant (*ire1*Δ), and *hxl1* gene-deficient mutant (*hxl1*Δ) were cultured in Sabouraud dextrose liquid medium at 27°C, 37°C, or 40°C, and growth was monitored by measuring absorbance at 630 nm. (**C**) The parent strain (Parent), *ire1* gene-deficient mutant (*ire1*Δ), and *hxl1* gene-deficient mutant (*hxl1*Δ) were cultured om SA medium at 27°C for 2 days. *T. asahii* cells were suspended in saline and serially diluted tenfold. Aliquots (5 µL) were spotted onto SA medium containing dithiothreitol (DTT; 15 mM), H_2_O_2_ (4 mM), Congo red (300 µg/mL), and sodium dodecyl sulfate (SDS; 0.01%), and incubated at 27°C for 72 h. (**D**) The *T. asahii* parent strain (Parent: 3.1 x 10^4^ cells/larva), *ire1* gene-deficient mutant (*ire1*Δ: 5.3 x 10^4^ cells/larva), and *hxl1* gene-deficient mutant (*hxl1*Δ:4.5 x 10^4^ cells/larva) were injected into silkworm hemolymph, and silkworm survival was monitored for 96 h at 37°C. Differences between the parent strain and mutant groups were assessed using the log-rank test based on Kaplan-Meier survival curves. *: P < 0.05. n = 10/group. (**E**) Dose-response analysis of *T. asahii* infection. The number of surviving silkworms at 37°C was determined 72 h after injection. Survival status was recorded as 1 (alive) or 0 (dead). n = 36-48/group. *: P < 0.05.

### Gene expression regulated by Ire1 and Hxl1 in *T. asahii*

In *C. neoformans*, expression of *SOD2*, which is involved in reactive oxygen species scavenging, is increased in both the *ire1* and *hxl1* gene-deficient mutants (15). Conversely, expression of *OST1*, *KAR2*, *ALG7*, and *DER1* is decreased in these mutants (15). Therefore, we investigated whether Ire1 and Hxl1 regulate the expression of *SOD1-3*, *CAT1-2*, *OST1*, *KAR2*, *ALG7*, and *DER1* in *T. asahii.* In *T. asahii*, the expression levels of *CAT2*, *SOD1*, and *SOD2* were reduced in the *ire1-* and *hxl1* gene-deficient mutants compared with the parent strain (Fig. 12A). In contrast, expression of *OST1*, *KAR2*, *ALG7*, and *DER1* was unchanged in these mutants (Fig. 12B). These findings suggest that Ire1 and Hxl1 regulate the expression of *CAT2*, *SOD1*, and *SOD2* in *T. asahii*.

**Fig. 12.**
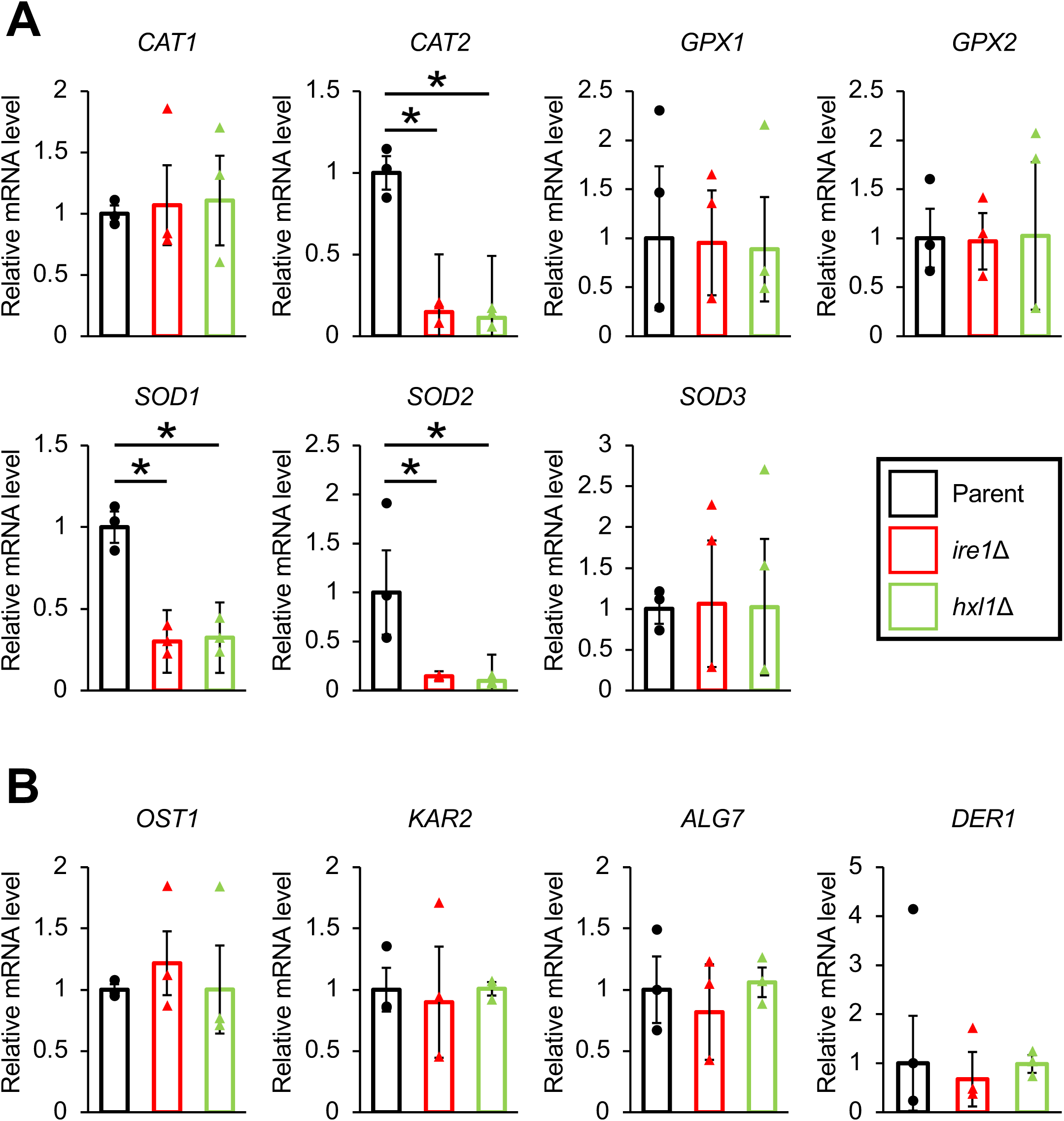
Decreased expression of antioxidant-related genes in *ire1* gene-deficient and *hxl1* gene-deficient *T. asahii* mutants. The parent strain (Parent), *ire1* gene-deficient mutant (*ire1*Δ), and *hxl1* gene-deficient mutant (*hxl1*Δ) were cultured on SA medium at 27°C for 2 days, followed by incubation in Sabouraud dextrose liquid medium containing DTT (3 mM) for 30 min. Gene expression levels were measured by quantitative RT-PCR. (**A**) mRNA levels of reactive oxygen species genes (*CAT1*, *CAT2*, *GPX1*, *GPX2*, *SOD1*, *SOD2*, and *SOD3)* were normalized to *ACT1* and expressed relative to the parent strain. (**B**) mRNA levels of UPR-related genes (*OST1*, *KAR2*, *ALG7*, and *DER1)* were normalized to *ACT1* and expressed relative to the parent strain. n = 3/group. *: P < 0.05.

## Discussion

In this study, we found that DTT treatment induces *hxl1* mRNA splicing in *T. asahii* and that *ire1* is required for this process. Furthermore, we demonstrated that *ire1* and *hxl1* play important roles in normal growth, tolerance to high-temperature stress, ER stress, oxidative stress, and cell wall and membrane stress, as well as in virulence in the silkworm infection model. These findings indicate that Ire1-mediated *hxl1* mRNA splicing regulates growth, stress resistance, and virulence in *T. asahii*.

The *hxl1* mRNA of *T. asahii* was spliced in response to DTT, an inducer of ER stress. In *C. neoformans*, Ire1 mediates *hxl1* mRNA splicing in response to DTT-induced stress (15). Previous *in silico* analyses suggested that *T. asahii hxl1* mRNA might be spliced in a manner similar to that in *C. neoformans hxl1* (21). In addition, a gene encoding a protein highly homologous to *C. neoformans* Ire1 (XP_012047375.1) was identified in the *T. asahii* genome. Domain analysis revealed that this protein contains the Luminal_IRE1 domain, which functions as a UPR sensor; the STKc_IRE1 domain, involved in autophosphorylation; and the RNase_Ire1 domain, responsible for splicing target mRNAs. Based on these features, we inferred that XP_014178218.1 encodes the Ire1 protein in *T. asahii*. In *ire1* gene-deficient mutants, DTT did not induce *hxl1* mRNA splicing. Taken together, we concluded that ER stress tolerance in *T. asahii* is mediated by an Ire1-dependent *hxl1* mRNA splicing mechanism, similar to that in *C. neoformans*.

The *T. asahii ire1* gene-deficient mutants were sensitive to DTT, H₂O₂, Congo Red, and SDS. The *ire1* gene-deficient mutants of *C. neoformans* and *C. albicans* show impaired ER stress resistance (15, 17). Consistent with this, *T. asahii ire1* gene-deficient mutants exhibited delayed growth on SA medium containing DTT. In contrast, their growth was not delayed on SA medium containing TM. DTT induces ER stress by non-specifically disrupting protein disulfide bonds within the ER as a reducing agent, whereas TM induces ER stress by inhibiting glycosyltransferases involved in protein glycosylation (33, 34). These results suggest that the differential sensitivity of *T. asahii ire1* gene-deficient mutants may be due to the distinct mechanisms of action of DTT and TM. Ire1 in *C. neoformans* contributes to oxidative stress resistance (15). Consistent with this, *T. asahii ire1* gene-deficient mutants delayed growth on SA medium containing H₂O₂ compared with the parent strain, suggesting that Ire1 is involved in oxidative stress tolerance in *T. asahii*. Ire1 in *C. neoformans* and *C. albicans* also contributes to cell wall stress resistance (15, 17, 31). Similarly, *T. asahii ire1* gene-deficient mutants exhibited delayed growth on SA medium containing Congo Red, suggesting a role in cell wall stress tolerance. In addition, Ire1 contributes to cell membrane stress resistance in these fungi (16, 17, 30), and *T. asahii ire1* gene-deficient mutants exhibited delayed growth on SA medium containing SDS, indicating involvement in cell membrane stress tolerance (16, 17, 31). Elucidating the relationship between Ire1 and tolerance to oxidative stress, as well as to cell wall and membrane stress in *T. asahii* remains an important area for future research. In *C. neoformans*, Ire1 is involved in pathogenicity in *G. mellonella* and mice, and it also contributes to the pathogenicity in *C. deuterogattii*, *C. albicans*, and *A. fumigatus* in mice (14–17). In this study, silkworms infected with *T. asahii ire1* gene-deficient mutants survived for 96 h after infection. These results suggest that Ire1 is involved in the stress tolerance and virulence of *T. asahii*.

The Hxl1 protein of *T. asahii*, like that of other pathogenic fungi, possesses a bZIP domain, which functions as a DNA-binding domain. Because DTT induces Ire1-dependent *T. asahii hxl1* mRNA splicing, we hypothesized that Hxl1 acts as a downstream transcription factor of Ire1 to regulate target gene expression. In *C. neoformans, hxl1* gene-deficient mutants are sensitive to ER stress induced by DTT and other compounds (15). Similarly, *T. asahii hxl1* gene-deficient mutants were sensitive to DTT, suggesting that Ire1-mediated *hxl1* mRNA splicing contributes to ER stress in *T. asahii*. Moreover, Hxl1 in *C. neoformans* contributes to oxidative stress resistance (15), and the *T. asahii hxl1* gene-deficient mutant exhibited sensitivity to H₂O₂. In addition, Hxl1 contributes to cell wall stress resistance in *C. neoformans*, and the *T. asahii hxl1* gene-deficient mutant exhibited delayed growth on SA medium containing Congo Red (15). These results suggest that Hxl1 is involved in oxidative and cell wall stress tolerance in *T. asahii*. In *C. neoformans*, Hxl1 also contributes to pathogenicity in *G. mellonella* and mice (15), and in *C. deuterogattii*, Hxl1 is involved in pathogenicity in mice (16). In the present study, silkworms infected with *T. asahii hxl1* gene-deficient mutants survived 96 h after infection, indicating reduced virulence. Therefore, these findings suggest that Hxl1 is involved in stress tolerance and virulence in *T. asahii*.

The phenotypes of the *T. asahii ire1* and *hxl1* gene-deficient mutants were compared. The *hxl1* gene-deficient mutant exhibited sensitivity to DTT and Congo red comparable to that of the *ire1* gene-deficient mutant, and its virulence against silkworms was reduced. These findings suggest that Ire1 in *T. asahii* is involved in ER stress tolerance, cell wall stress tolerance, and virulence via Hxl1. The *hxl1* gene-deficient mutant also exhibited sensitivity to H₂O₂ and SDS, although to a lesser extent than the *ire1* gene-deficient mutant. Based on these findings, we hypothesized that Ire1 in *T. asahii* may contribute to resistance to H₂O₂ and SDS through both Hxl1-dependent and non-Hxl1-dependent pathways. Splicing of *hxl1* mRNA by Ire1 in *T. asahii* affected the expression of genes involved in antioxidant activity. In *ire1* and *hxl1* gene-deficient mutants, the expression levels of *CAT2*, *SOD1*, and *SOD2* were reduced under DTT conditions. Therefore, we propose that Ire1-induced *hxl1* mRNA splicing contributes to stress tolerance in *T. asahii* by regulating the expression of enzymes involved in reactive oxygen species degradation.

A limitation of this study is that we were unable to establish a complemented strain by reintroducing the *ire1* gene into the *ire1* gene-deficient mutant. This was due to the large size of the *ire1* gene, which prevented construction of a suitable targeting DNA fragment. Therefore, we independently established three *ire1* gene-deficient mutants and confirmed that their phenotypes were consistent. In addition, although infection experiments were performed using silkworms, an invertebrate model, no experiments were performed in mammalian models. Determining whether *ire1* and *hxl1* in *T. asahii* are involved in virulence in mice will be an important topic for future research. In addition, identifying the DNA sequences targeted by Hxl1 remains a challenge for future studies.

In conclusion, we demonstrated that DTT induces Ire1-dependent *hxl1* mRNA splicing in *T. asahii*, and that Ire1 and Hxl1 play crucial roles in normal growth; tolerance to heat, ER, oxidative, cell wall and membrane stress; and virulence. Furthermore, we showed that Ire1 and Hxl1 influence the expression of genes involved in antioxidant defense. These findings suggest that Ire1 and Hxl1 may serve as potential therapeutic targets for the treatment of *T. asahii* infections.

## Acknowledgments

We thank Renta Endo, Momoka Matsumura, and Tomoya Sanbongi (Meiji Pharmaceutical University) for technical assistance rearing the silkworms. We thank Yuna Tabata and Yasuko Yamaguchi (Meiji Pharmaceutical University) for helping to generate the gene-deficient mutants. We also thank SciTechEdit International LLC (Highlands Ranch, CO, USA) for English language editing. This study was supported by JSPS KAKENHI grant number JP23K06141 and JP26K09636 (Scientific Research (C) to Y.M.), in part by the Research Program on Emerging and Re-emerging Infectious Diseases of the Japan Agency for Medical Research and Development, AMED (Grant number JP24fk0108679h0402 to T.S.), and the Research and Implementation Promotion Program through open innovation grants(Grant# JPJ011937, to the consortium where Y.M. serves as a member) from the Project of the Bio-oriented Technology Research Advancement Institution (BRAIN).

